# ERCC1 mice, unlike other premature aging models, display accelerated epigenetic age

**DOI:** 10.1101/2022.12.28.522011

**Authors:** Kevin Perez, Alberto Parras, Cheyenne Rechsteiner, Amin Haghani, Robert Brooke, Calida Mrabti, Lucas Schonfeldt, Steve Horvath, Alejandro Ocampo

## Abstract

Over the last decades, several premature aging mouse models have been developed to study aging and identify interventions that can delay age-related diseases. Yet, it is still unclear whether these models truly recapitulate natural aging. Here, we analyzed DNA methylation in multiple tissues of four previously reported mouse models of premature aging (ERCC1, LAKI, POLG and XPG). We estimated DNA methylation (DNAm) age of these samples using the Horvath clock. The most pronounced increase in DNAm age could be observed in ERCC1 mice, a strain which exhibits a deficit in DNA nucleotide excision repair. In line with these results, we detected an increase in epigenetic age in fibroblasts isolated from patients with progeroid syndromes associated with mutations in DNA excision repair genes. These findings highlight ERCC1 as a particularly attractive mouse model to study aging in mammals and suggest a strong connection between DNA damage and epigenetic dysregulation during aging.

## MAIN TEXT

The world’s population is growing older. Since aging represents the strongest risk factor for most human diseases, it is therefore key to identify anti-aging interventions that could delay or even reverse the aging process^1^. Towards this goal, several accelerated aging mouse models have been developed to study the aging process^2,3^, some of them stemming from existing human disorders^4,5^. In this line, premature aging rodents could speed up the discovery of anti-aging interventions by shortening the experimental time, but only if the results can be translatable to natural aging. Nevertheless, the physiological relevance of these models and whether they truly recapitulate or phenocopy natural aging remains controversial. Epigenetic changes are one of several hallmarks of aging in numerous organisms^6^. The importance of epigenetic changes in mammals has been reinforced by the development of epigenetic clocks that can accurately estimate age in multiple tissues and all mammalian species^7-11^. Interestingly, several anti-aging interventions have been shown to reverse these clocks^12^, including cellular reprogramming^13-16^. Here, we sought to assess the relevance of several premature aging mouse models to study aging. Toward this end, we analyzed mouse models of segmental progeria by assessing the epigenetic age of multiple tissues and organs using epigenetic clocks based on DNA methylation.

Specifically, we analyzed the epigenetic age (“Horvath Pan Tissue clock”)^17^ of five tissues of four commonly used premature aging models including: ERCC1, XPG, LAKI and POLG mice. These mouse strains cause premature aging through various biological mechanisms by carrying mutations that lead to the manipulation of different hallmarks of aging. Specifically, ERCC1^18^ and XPG^19^ mice exhibit a deficit in nucleotide excision repair (NER) of the nuclear DNA, POLG mice show accumulation of mitochondrial DNA mutations^20,21^ and lastly LMNA knock-in (LAKI) mice suffer nuclear lamina defects^22,23^. To perform comparative studies in these strains, we assessed the DNA methylation age (DNAm) in ERCC1^KO/Δ^, XPG^KO/KO^, LAKI^TG/TG^ and POLG^TG/TG^ mice at several timepoints including during post-natal development, at median survival, and in old age, relative to each model’s own lifespan. Both proliferative (blood and skin) and more terminally differentiated tissues (liver, cerebral cortex, and skeletal muscle) were analyzed at these ages (Figure 1a). During the generation of experimental mice, we noticed that while LAKI^TG/TG^ and POLG^TG/TG^ mice were born at a predicted Mendelian frequency, ERCC1^KO/Δ^ and XPG^KO/KO^ showed a perinatal lethality (Figure S1a). Furthermore, as previously reported the four premature aging animals were significantly smaller and exhibited reduced body weight compared to their control littermates as expected (Figure 1b). Before analyzing the progeria models, we first looked at the clock performance in the control littermate WT mice (C57BL6J and C57BL6J|FVB hybrid backgrounds), a quality check that methylation can accurately predict chronological age in multiple tissues. The chronological age prediction in these two different backgrounds was highly accurate in blood (C57BL6J, RMSE: 2.08wk, r = 0.99; C57BL6J|FVB, RMSE: 2.55wk, r = 0.95) and provided sufficient accuracy in the other tissues (Figure S1b and Table S1), confirming the precision of the DNAm clock to predict age, particularly in blood. Next, we determined the DNAm age in the five tissues of ERCC1^KO/Δ^, LAKI^TG/TG^ and XPG^KO/KO^ at 8 weeks, and POLG^TG/TG^ at 30 weeks of age corresponding to the relative median survival of the strain. Strikingly, ERCC1^KO/Δ^ was the only premature aging model where we observed increased biological age compared to control littermates (Figure 1c). Importantly, the biological age of ERCC1^KO/Δ^ mice was most increased in blood [WT: 6.85w (1.62), KO/Δ: 12.46w (1.08)], but was also significantly increased in brain, liver, skeletal muscle and skin, tissues and organs known to be affected in this mouse model. Conversely, we did not detect any acceleration in DNAm age at 8 weeks in LAKI or XPG mice, nor in POLG mice at 30 weeks in any tissue (Figure 1c). This result indicates that only ERCC1 aging mouse model shows a significant increase in epigenetic age at the median lifespan.

**FIGURE 1.**
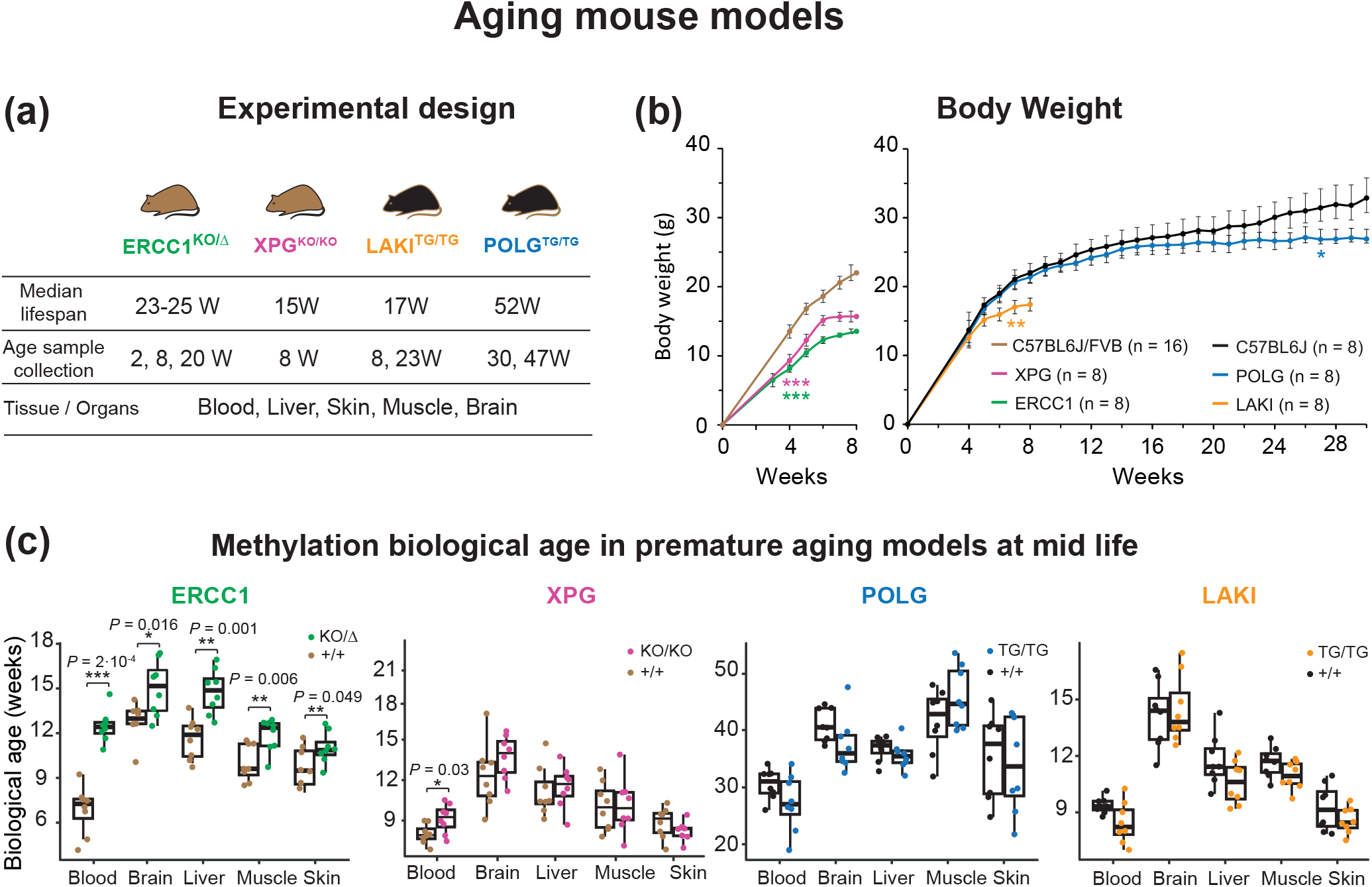
DNA methylation in premature aging mouse models. **(a)** Schematic representation of premature mouse strains and littermate controls, tissues collected, and timepoints taken. **(b)** Evolution of body weight (grams) of mutant and controls mice from 4 weeks until the euthanize point, data are mean ± SEM. **(c)** Methylation biological age (in weeks) of ERCC1^KO/Δ^, XPG^KO/KO^, LAKI^TG/TG^ at 8 weeks and POLG^TG/TG^ at 30 weeks. Data are represented as box plots (center line shows median, box shows 25th and 75th percentiles and whiskers show minimum and maximum values and statistical significance was assessed by two-sided unpaired t-test.

Subsequently, and with the goal of confirming this observation, we analyzed the methylation age at different times points during the lifespan of the mice including, ERCC1^KO/Δ^ (2, 8 and 20 weeks), LAKI^TG/TG^ (8 and 23 weeks), and POLG^TG/TG^ (30 and 47 weeks). Interestingly, in the ERCC1^KO/Δ^ mice, biological age was increased mildly at 2 weeks old in blood, but not in other tissues. However, at 20 weeks, DNAm age was significantly accelerated in blood, liver, and skin (Figure 2a and Table S2). Conversely, as we observed at earlier timepoints, DNAm age was not changed in any of the analyzed tissues at 14.4 weeks in LAKI^TG/TG^ mice, nor in POLG^TG/TG^ mice at 47 weeks (Figure S1c). Together, our results further confirm that the biological age measured by DNA methylation is increased only in the ERCC1 mouse model of premature aging, at multiple ages, with blood being the tissue with the strongest statistical power. Importantly, when the same analysis was restricted to either male or female only, the same trend appeared, with increased DNAm age primarily in the ERCC1 mouse model. Next, we wondered whether the observed differences between methylation age and chronological age in ERCC1 mice were constant or changed throughout life. To determine this accelerated aging rate, we calculated the slope between biological and chronological age in each tissue in ERCC1 +/+ vs. KO/Δ mice. Importantly, the rate was significantly different in blood (Slope: WT = 0.78, KO/Δ = 1.29), skeletal muscle (Slope: WT = 0.84, KO/Δ = 1.17) and brain (Slope: WT = 0.91, KO/Δ = 1.2) (Figure 2b), demonstrating that the difference between biological and chronological age increased during life in ERCC1^KO/Δ^ mice.

**FIGURE 2.**
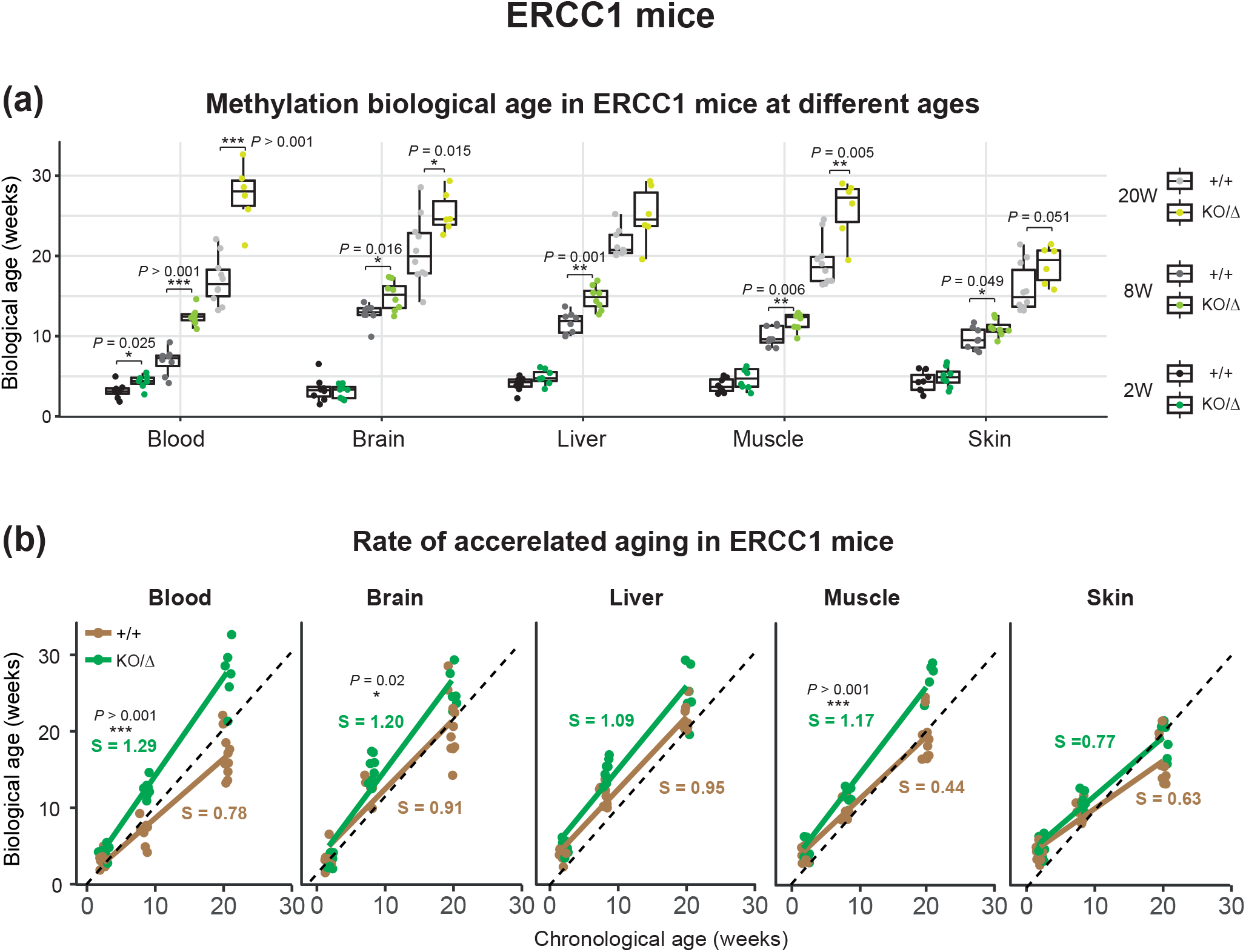
DNA methylation ERCC1 mice. **(a)** Methylation biological age (in weeks) of ERCC1^KO/Δ^ mice at 2, 8 and 20 weeks in multiple organs/tissues and WT littermate controls estimated by Horvath clock. Data are represented as box plots (center line shows median, box shows 25th and 75th percentiles and whiskers show minimum and maximum values and statistical significance was assessed by two-sided unpaired t-test. **(b)** Slope of aging in ERCC1^KO/Δ^ and controls mice in tissues analyzed from 2 to 20 weeks old. Significance of the interaction term in the linear regression was analyzed.

Lastly, to investigate the potential relevance of these findings to human patients, we analyzed the DNAm age of samples obtained from patients affected by diseases caused by mutations in DNA excision repair genes associated with aging phenotypes: Xeroderma Pigmentosum (XP) affecting *ERCC5*^24^, and Cockayne Syndrome (CS) type A (CSA) affecting *ERCC8* and type B (CSB) affecting *ERCC6*^25^. Towards this goal, we profiled DNAm age from fibroblasts derived from patients at multiple ages: control (1, 5, 11-year-old), CSA (1, 3, 5-year-old), CSB (3, 8, 10-year-old), XP (1, 2, 5-year-old). For this analysis only, we selected the DNAm age from the “Skin&Blood” Clock, as this has previously been shown to be more accurate than the “PanTissue” clock to assess age of human fibroblasts^26^, a finding that we also confirmed in our own dataset (Figure 3a). Importantly, the DNAm age was significantly higher in the affected patients compared to control samples (Figure 3a). Finally, we calculated the difference between DNAm age and chronological age for each sample, detecting a significant increase for the XP patients and a strong tendency in the rest of the disease samples (Figure 3b). Overall, these results indicate that human progeroid syndromes associated with mutations in DNA excision repair genes display accelerated epigenetic age.

**FIGURE 3.**
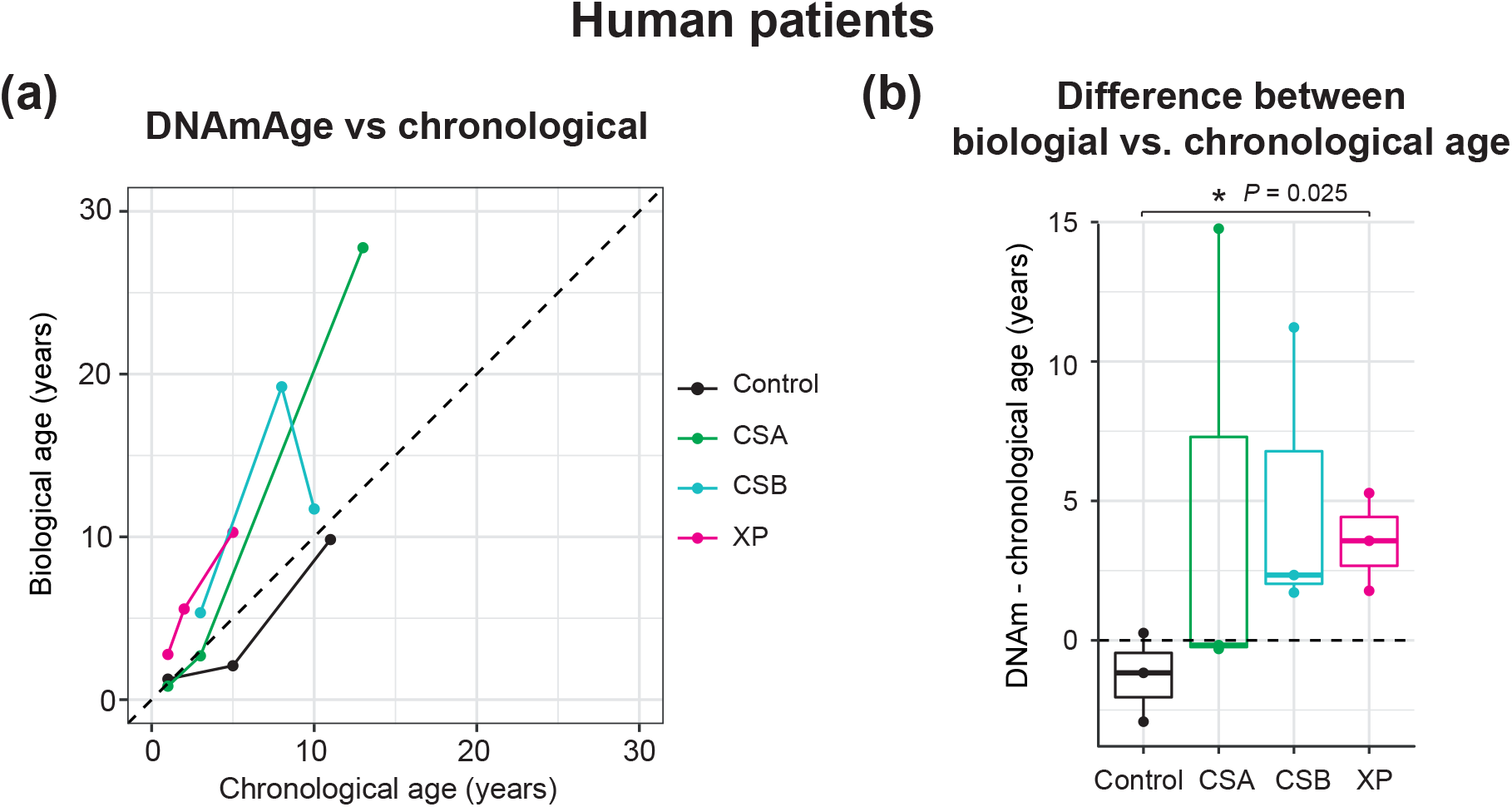
DNA methylation in fibroblasts from human premature aging diseases. **(a)** DNAm age versus chronological age (in years) and **(b)** difference between biological and chronological age in human samples in fibroblasts isolated from individual with Cockayne Syndrome A (CSA), Cockayne Syndrome B (CSB), Xeroderma Pigmentosum (XP) and controls analyzed by Skin&Blood Clock. Data are represented as box plots (center line shows median, box shows 25th and 75th percentiles and whiskers show minimum and maximum values and statistical significance was assessed by two-sided unpaired t-test.

Although premature aging models have been widely used to study aging and evaluate antiaging interventions, their physiological relevance for the study of aging has not been deeply investigated. Here, we analyzed the biological age (“Horvath clock”) of four premature aging mouse models (ERCC1, POLG, XPG, LAKI) and demonstrated that only ERCC1 mice truly shows accelerated aging.

Depletion of ERCC1 protein results in a defect in DNA repair, leading to an accumulation of DNA mutations in multiple tissues and organs. Importantly, DNA damage has been proposed as one of the most central hallmarks of aging, as well as a causative driver^27,28^. Here, we show that a defective DNA repair mechanism leads to epigenetic aging, strongly suggesting a link between DNA damage and epigenetic dysregulation. Interestingly, dietary restriction, the most robust anti-aging intervention, dramatically extends lifespan of ERCC1^KO/Δ^ mice^29^ and knocking down of *ERCC1* gene in blood specifically causes premature aging^30^. Furthermore, we noted that even though DNAm age was increased in ERCC1 mice already at 2 weeks, greater changes were observed in older animals indicating a progressive age acceleration during aging. In this line, we postulate that a higher DNA repair capacity during development^31^ or embryonic reprogramming programs, which might prevent potential epigenetic dysregulation as consequence of DNA damage, could protect the animals during gestation. Taken together, these results suggest that ERCC1 mice stand perhaps as one of the most relevant mouse models of premature aging.

The methylation clock was more accurate in blood, a rapidly proliferative tissue that undergoes constant regeneration, in which the most significant and strongest differences between ERCC1 and control mice were observed. Therefore, due to its easy collection and strong sensitivity for epigenetic aging, we propose the use of blood as one of the best choices to study and analyze the effect of anti-aging interventions. Lastly, although multiple groups have examined the biological age of human diseases associated with premature aging, no changes in DNAm age have been observed in the blood of Hutchinson-Gilford progeria syndrome patients^32^. On the other hand, a significant increase in biological age was seen in samples from Werner^33^, Down syndrome even in newborns^34^ in several human overgrowth syndromes including Sotos syndrome^35^ and Tatton-Brown-Rahman syndrome^36^ and very recently in Leigh Syndrome and mitochondrial encephalopathy with lactic acidosis and stroke-like episodes (MELAS) patients^37^. Other studies have identified changes in DNAm in premature aging models, independent of the DNA methylation clocks^38-40^. Our survey of mouse models of premature aging may be expanded to alternative premature aging models^2^, or additional tissues and timepoints. Likewise, it would be interesting to also assess biological age using newly developed clocks, such as transcriptomic, proteomic or chromatin accessibility clocks^41-43^.

## ACKNOWLEDGMENTS

We would like to thank all members of the Ocampo laboratory for helpful discussions. We would like to thank the teams of mouse facilities at the University of Lausanne including Francis Derouet and Lisa Arlandi (animal facility at Epalinges), and Laurent Lecomte (animal facility of the Department of Biomedical Sciences) and colleagues. We thank Prof. Jan Hoeijmakers and Prof. Ingrid van der Pluijm for providing us the ERCC1 and XPG mice, and Prof. Carlos Lopez Otin and Prof. Andrea Ablasser for providing us the LAKI mice.

This work was supported the Swiss National Science Foundation (SNSF), the University of Lausanne, and the Canton Vaud.

## AUTHOR CONTRIBUTIONS

K.P. performed data and statistical analysis. A.P. generated mouse strains and collected tissues. C.M and L.S. were involved in culture and DNA extraction from human cells. C.R. extracted DNA from mice. S.H., A.H. analyzed data and made a critically revision. A.O. directed, supervised the study, designed the experiments, and reviewed the manuscript. K.P. and A.P generated the figures and wrote the manuscript with input from all authors.

## DATA AVAILABILITY STATEMENT

The data supporting the findings of this study are available from the corresponding author upon reasonable request.

The mammalian methylation array is available from the nonprofit Epigenetic Clock Development Foundation (https://clockfoundation.org/)

## EXPERIMENTAL PROCEDURES

### Animal housing

All the experimental experiment were performed in accordance with Swiss legislation after the approval from the local authorities (Cantonal veterinary office, Canton de Vaud, Switzerland). Mice were housed in groups of five per cage with a 12hr light/dark cycle between 06:00 and 18:00 in a temperature-controlled environment at 25°C and humidity between 40 % and 70 %, with free access to water and food. Wild type (WT) and premature aging mouse models used in this study were generated by breeding (Figure S1a) and housed together until they reached the desired age in the Animal Facilities of Epalinges and Department of Biomedical Science of the University of Lausanne.

### Mouse strains

ERCC1^KO/Δ 44^ and XPG^KO/KO^ mice^19^ and littermate controls (ERCC1^+/+^ and XPG^+/+^) were used in C57BL6J|FVB hybrid background. POLG^D257A/D257A^, herein referred to as POLG^TG/TG 20,21^ and LAKI^TG/TG 22^ and sibling controls (POLG^+/+^ and LAKI^+/+^) were generated in C57BL6J background.

### Mouse monitoring and euthanasia

All mice were monitored at least three times per week to evaluate their activity, posture, alertness, body weight, presence of tumors or wound, and surface temperature. Males and females were euthanized at the specific timepoints by CO_2_ inhalation (6 min, flow rate 20% volume/min). Subsequently, before perfusing the mice with saline, blood was collected from the heart. Finally, multiple organs and tissues were collected in liquid nitrogen and used for DNA extraction to perform MethylArray.

### Cell culture and maintenance

Human fibroblasts were obtained from the Coriell cell repositories and cultured in DMEM (Gibco, 11960085) with 10% FBS (Hyclone, SH30088.03) containing non-essential amino acids, GlutaMax and Sodium Pyruvate (Gibco, 11140035, 35050061, 11360039) at 37°C in hypoxic conditions (3% O_2_). Subsequently, fibroblasts were passaged and cultured according to standard protocols.

### DNA extractions

Total DNA was extracted from tissues and cells using Monarch Genomic DNA Purification Kit (New England Biolab, T3010L) and protocols were carefully followed. Tissues were cut into small pieces to ensure rapid lysis. Total DNA concentrations were determined using the Qubit DNA BR Assay Kit (Thermofisher, Q10211).

### DNA methylation clock

The mouse clock was developed in^17^. We used the “Pan Tissue” mouse clock since we analyzed different tissues. The software code of the mouse clocks can be found in the supplements of^17^.

The mouse methylation data were generated on the small and the extended version of HorvathMammalMethylChip^45^. We used the SeSaMe normalization method^46^. Human methylation data were generated on the Illumina EPIC array platforms that profiles 866k cytosines. We used the noob normalization method implemented in the R function preprocessNoob. The human DNAm age was estimated using the Skin&blood clock algorithm^26^.

### Statistical analysis

Unsupervised hierarchical clustering based on interarray correlation coefficients was used to identify putative technical outliers. One liver sample with negative methylation age was removed. All plots were generated using the R software package ggplot2. Statistical differences between groups were assessed using a two-tailed unpaired Student’s t-test. Clock performance was assessed by correlation (Pearson coefficient) and Random Mean Square Error (RMSE), using the R software. To determine if there was a significant difference in the slope of aging between WT and transgenic mice, we looked at the significance of the interaction term in the linear regression: DNAm age ∼ WT/TG + Age + WT/TG*Age.

---

**SUPPLEMENTARY FIGURE 1.**
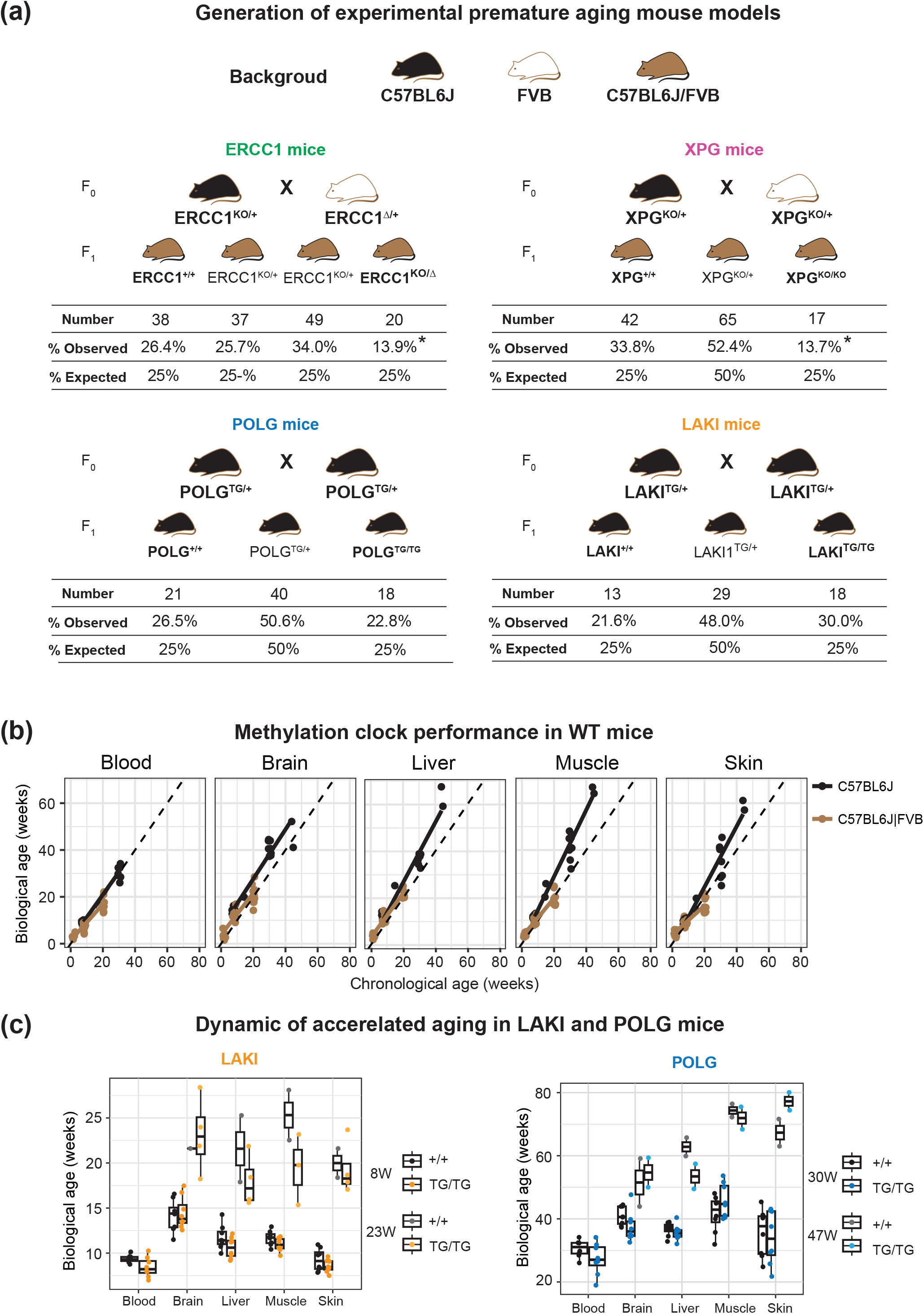
DNA methylation in premature aging mouse models additional data. **(a)** Breeding protocol to generate the four premature mouse strains and littermate control mice. Statistical significance was assessed by Pearson’s chi-squared test. **(b)** Correlation between biological and chronological age (in weeks) in WT control mice in C57BL6J and C57BL6J|FVB backgrounds in analyzed tissues from 2-to 47-week-old. **(c)** Methylation biological age of POLG^TG/TG^ (at 30 and 47 weeks old) and LAKI^TG/TG^ (at 8 and 23 weeks) in multiple organs/tissues and WT littermate controls by Horvath clock. Data are represented as box plots (center line shows median, box shows 25th and 75th percentiles and whiskers show minimum and maximum values) and statistical significance was assessed by two-sided unpaired t-test.

**Table S1.**

| Tissue | C57BL6J [RMSE (r)] | C57BL6J \| FVB [RMSE (r)] |
| --- | --- | --- |
| Blood | 2.08 (0.99) | 2.55 (0.95) |
| Brain | 8.71 (0.96) | 4.13 (0.89) |
| Liver | 8.49 (0.98) | 3.04 (0.97) |
| Muscle | 11.2 (0.98) | 2.51 (0.95) |
| Skin | 7.59 (0.96) | 3.21 (0.91) |

**Table S2.**
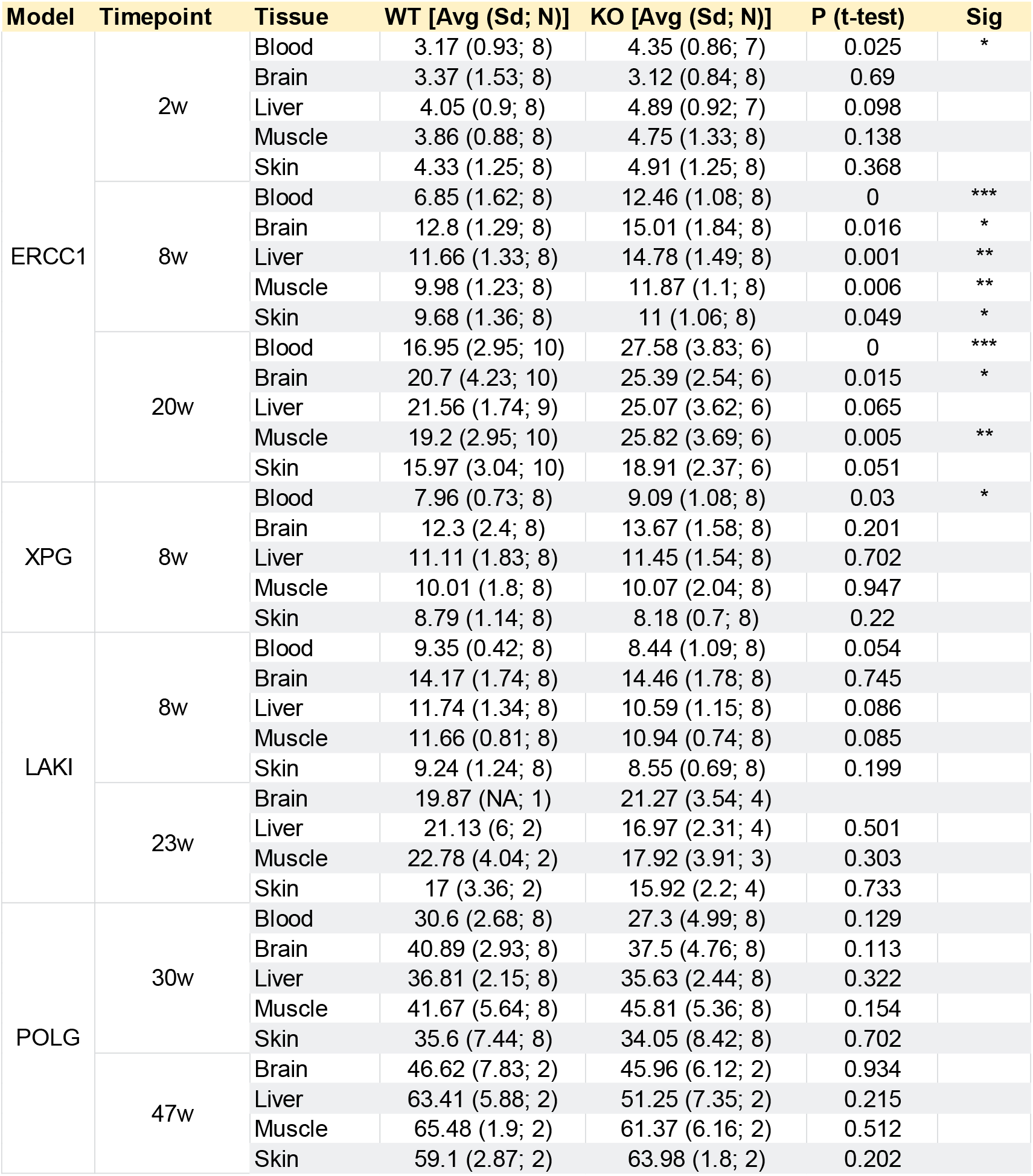

## REFERENCES

1 Partridge, L., Deelen, J. & Slagboom, P. E. Facing up to the global challenges of ageing. Nature 561, 45–56, doi:10.1038/s41586-018-0457-8 (2018).

2 Koks, S. et al. Mouse models of ageing and their relevance to disease. Mech Ageing Dev 160, 41–53, doi:10.1016/j.mad.2016.10.001 (2016).

3 Liao, C. Y. & Kennedy, B. K. Mouse models and aging: longevity and progeria. Curr Top Dev Biol 109, 249–285, doi:10.1016/B978-0-12-397920-9.00003-2 (2014).

4 Bohr, V. A. Human premature aging syndromes and genomic instability. Mech Ageing Dev 123, 987–993, doi:10.1016/s0047-6374(02)00039-8 (2002).

5 Brosh, R. M., Jr. & Bohr, V. A. Human premature aging, DNA repair and RecQ helicases. Nucleic Acids Res 35, 7527–7544, doi:10.1093/nar/gkm1008 (2007).

6 Lopez-Otin, C., Blasco, M. A., Partridge, L., Serrano, M. & Kroemer, G. The hallmarks of aging. Cell 153, 1194–1217, doi:10.1016/j.cell.2013.05.039 (2013).

7 Horvath, S. DNA methylation age of human tissues and cell types. Genome Biol 14, R115, doi:10.1186/gb-2013-14-10-r115 (2013).

8 Simpson, D. J. & Chandra, T. Epigenetic age prediction. Aging Cell 20, e13452, doi:10.1111/acel.13452 (2021).

9 Ake Lu, V. P., Fei, Z., Raj, K. & Horvath, S. Universal DNA Methylation Age Across Mammalian Tissues. Innovation in Aging 5, 410–410, doi:10.1093/geroni/igab046.1588 (2021).

10 Bell, C. G. et al. DNA methylation aging clocks: challenges and recommendations. Genome Biol 20, 249, doi:10.1186/s13059-019-1824-y (2019).

11 Bergsma, T. & Rogaeva, E. DNA Methylation Clocks and Their Predictive Capacity for Aging Phenotypes and Healthspan. Neurosci Insights 15, 2633105520942221, doi:10.1177/2633105520942221 (2020).

12 Field, A. E. et al. DNA Methylation Clocks in Aging: Categories, Causes, and Consequences. Mol Cell 71, 882–895, doi:10.1016/j.molcel.2018.08.008 (2018).

13 Simpson, D. J., Olova, N. N. & Chandra, T. Cellular reprogramming and epigenetic rejuvenation. Clin Epigenetics 13, 170, doi:10.1186/s13148-021-01158-7 (2021).

14 Sarkar, T. J. et al. Transient non-integrative expression of nuclear reprogramming factors promotes multifaceted amelioration of aging in human cells. Nat Commun 11, 1545, doi:10.1038/s41467-020-15174-3 (2020).

15 Browder, K. C. et al. In vivo partial reprogramming alters age-associated molecular changes during physiological aging in mice. Nature Aging, doi:10.1038/s43587-022-00183-2 (2022).

16 Lu, Y. et al. Reprogramming to recover youthful epigenetic information and restore vision. Nature 588, 124–129, doi:10.1038/s41586-020-2975-4 (2020).

17 Mozhui, K. et al. Genetic loci and metabolic states associated with murine epigenetic aging. Elife 11, doi:10.7554/eLife.75244 (2022).

18 Weeda, G. et al. Disruption of mouse ERCC1 results in a novel repair syndrome with growth failure, nuclear abnormalities and senescence. Curr Biol 7, 427–439, doi:10.1016/s0960-9822(06)00190-4 (1997).

19 Barnhoorn, S. et al. Cell-autonomous progeroid changes in conditional mouse models for repair endonuclease XPG deficiency. PLoS Genet 10, e1004686, doi:10.1371/journal.pgen.1004686 (2014).

20 Kujoth, G. C. et al. Mitochondrial DNA mutations, oxidative stress, and apoptosis in mammalian aging. Science 309, 481–484, doi:10.1126/science.1112125 (2005).

21 Trifunovic, A. et al. Premature ageing in mice expressing defective mitochondrial DNA polymerase. Nature 429, 417–423, doi:10.1038/nature02517 (2004).

22 Osorio, F. G. et al. Splicing-directed therapy in a new mouse model of human accelerated aging. Sci Transl Med 3, 106ra107, doi:10.1126/scitranslmed.3002847 (2011).

23 Varga, R. et al. Progressive vascular smooth muscle cell defects in a mouse model of Hutchinson-Gilford progeria syndrome. Proc Natl Acad Sci U S A 103, 3250–3255, doi:10.1073/pnas.0600012103 (2006).

24 Rizza, E. R. H. et al. Xeroderma Pigmentosum: A Model for Human Premature Aging. J Invest Dermatol 141, 976–984, doi:10.1016/j.jid.2020.11.012 (2021).

25 Laugel, V. in GeneReviews((R)) (eds M. P. Adam et al.) (1993).

26 Horvath, S. et al. Epigenetic clock for skin and blood cells applied to Hutchinson Gilford Progeria Syndrome and ex vivo studies. Aging (Albany NY) 10, 1758–1775, doi:10.18632/aging.101508 (2018).

27 Yousefzadeh, M. et al. DNA damage-how and why we age? Elife 10, doi:10.7554/eLife.62852 (2021).

28 Schumacher, B., Pothof, J., Vijg, J. & Hoeijmakers, J. H. J. The central role of DNA damage in the ageing process. Nature 592, 695–703, doi:10.1038/s41586-021-03307-7 (2021).

29 Vermeij, W. P. et al. Restricted diet delays accelerated ageing and genomic stress in DNA-repair-deficient mice. Nature 537, 427–431, doi:10.1038/nature19329 (2016).

30 Yousefzadeh, M. J. et al. An aged immune system drives senescence and ageing of solid organs. Nature 594, 100–105, doi:10.1038/s41586-021-03547-7 (2021).

31 Mitchell, D. L. & Hartman, P. S. The regulation of DNA repair during development. Bioessays 12, 74–79, doi:10.1002/bies.950120205 (1990).

32 Bejaoui, Y. et al. DNA methylation signatures in Blood DNA of Hutchinson-Gilford Progeria syndrome. Aging Cell 21, e13555, doi:10.1111/acel.13555 (2022).

33 Maierhofer, A. et al. Accelerated epigenetic aging in Werner syndrome. Aging (Albany NY) 9, 1143–1152, doi:10.18632/aging.101217 (2017).

34 Xu, K. et al. Accelerated epigenetic aging in newborns with Down syndrome. Aging Cell 21, e13652, doi:10.1111/acel.13652 (2022).

35 Martin-Herranz, D. E. et al. Screening for genes that accelerate the epigenetic aging clock in humans reveals a role for the H3K36 methyltransferase NSD1. Genome Biol 20, 146, doi:10.1186/s13059-019-1753-9 (2019).

36 Jeffries, A. R. et al. Growth disrupting mutations in epigenetic regulatory molecules are associated with abnormalities of epigenetic aging. Genome Res 29, 1057–1066, doi:10.1101/gr.243584.118 (2019).

37 Yu, T. et al. Premature aging is associated with higher levels of 8-oxoguanine and increased DNA damage in the Polg mutator mouse. Aging Cell 21, e13669, doi:10.1111/acel.13669 (2022).

38 Guastafierro, T. et al. Genome-wide DNA methylation analysis in blood cells from patients with Werner syndrome. Clin Epigenetics 9, 92, doi:10.1186/s13148-017-0389-4 (2017).

39 Heyn, H., Moran, S. & Esteller, M. Aberrant DNA methylation profiles in the premature aging disorders Hutchinson-Gilford Progeria and Werner syndrome. Epigenetics 8, 28–33, doi:10.4161/epi.23366 (2013).

40 Crochemore, C. et al. Epigenomic signature of the progeroid Cockayne syndrome exposes distinct and common features with physiological ageing. bioRxiv, 2021.2005.2023.445308, doi:10.1101/2021.05.23.445308 (2021).

41 Meyer, D. H. & Schumacher, B. BiT age: A transcriptome-based aging clock near the theoretical limit of accuracy. Aging Cell 20, e13320, doi:10.1111/acel.13320 (2021).

42 Lehallier, B., Shokhirev, M. N., Wyss-Coray, T. & Johnson, A. A. Data mining of human plasma proteins generates a multitude of highly predictive aging clocks that reflect different aspects of aging. Aging Cell 19, e13256, doi:10.1111/acel.13256 (2020).

43 Rechsteiner, C. et al. Development of a novel aging clock based on chromatin accessibility. bioRxiv, 2022.2008.2011.502778, doi:10.1101/2022.08.11.502778 (2022).

44 de Waard, M. C. et al. Age-related motor neuron degeneration in DNA repair-deficient Ercc1 mice. Acta Neuropathol 120, 461–475, doi:10.1007/s00401-010-0715-9 (2010).

45 Arneson, A. et al. A mammalian methylation array for profiling methylation levels at conserved sequences. Nat Commun 13, 783, doi:10.1038/s41467-022-28355-z (2022).

46 Zhou, W., Triche, T. J., Jr., Laird, P. W. & Shen, H. SeSAMe: reducing artifactual detection of DNA methylation by Infinium BeadChips in genomic deletions. Nucleic Acids Res 46, e123, doi:10.1093/nar/gky691 (2018).

